# Symmetrical Dimethylarginine as the Central Antigenic Determinant of Anti-Smith Autoantibodies in Systemic Lupus Erythematosus

**DOI:** 10.1101/2025.09.25.678492

**Authors:** Lars C. van Vliet, Annemarie L. Dorjée, Martijn van der Heide, Gerda M. Steup-Beekman, Thomas W.J. Huizinga, René E.M. Toes, Jolien Suurmond

**Author notes:** Corresponding author: Dr. J. Suurmond, Albinusdreef 2, LUMC - Postzone C1-R 2333 ZA, Leiden, The Netherlands.

## Abstract

**Objective:** Systemic lupus erythematosus (SLE) is a chronic autoimmune disease causing multi-organ damage. The most specific autoantibody response in SLE, present in 20-30% of patients, targets the Sm-protein and has been shown to recognize a linear Sm-derived B cell epitope containing a post-translational modification (PTM) of arginine termed symmetrical dimethyl arginine (sDMA). As autoantibodies to other PTM-modified proteins are often promiscuous, we aimed to determine the specificity and cross-reactivity of anti-Sm antibodies.

**Methods:** Specificity and promiscuity/cross-reactivity of anti-Sm IgG were measured by ELISA in SLE patients and healthy donors using peptides containing either sDMA or unmodified arginine. Inhibition and cross-reactivity were determined using competitive ELISA and affinity purification. Recognition of endogenous EBNA1 by anti-Sm IgG was performed by Western blot using lysates of EBV-bearing lymphoblastoid cell lines.

**Results:** The sDMA residue is recognized by anti-Sm+ SLE patients regardless of the peptide amino acid sequence, with a modest impact of amino acids flanking sDMA on recognition. Most notably, IgGs targeting sDMA comprise the overall majority (∼90%) of the anti-Sm antibody repertoire and are highly cross-reactive between SmD3_108-122_ and several sDMA-containing viral-derived epitopes, including full-length EBNA1.

**Conclusion:** Our data implicate that the majority of anti-Sm IgGs target the sDMA residue irrespective of its Sm-context, thus representing a prototypic anti-PTM response. These antibodies are highly promiscuous, recognizing several sDMA-modified targets, including naturally occurring viral sDMA-expressing epitopes. These findings suggest a new mechanism by which molecular mimicry of sDMA-modified viral proteins could contribute to a breach of tolerance in anti-Sm+ SLE patients.

**KEY MESSAGES:** *What is already known on this topic:*

- Symmetrical dimethylarginine (sDMA) residues are present on RGG/RG repeat regions within the SmD1, SmD3, and SmB/B⍰ subunits of the Sm protein complex *in vivo*.
- The introduction of sDMA is essential for recognition of minimal antigenic epitopes spanning amino acids #95–119 on the SmD1 and #108–122 on the SmD3 subunits by a specific subset of anti-Sm autoantibodies.

*What this study adds:*

- We show that autoantibodies targeting sDMA residues are common in anti-Sm+ SLE patients and represent the majority of the anti-Sm autoantibody repertoire.
- We demonstrate that anti-sDMA autoantibodies are highly cross-reactive and can also bind to sDMA residues on other targets, including several virus-derived epitopes, thus representing a prototypic anti-modified protein antibody (AMPA) response directed against post-translationally modified proteins through symmetrical dimethylation of arginine.

*How this study might affect research, practice, or policy:*

- The high cross-reactivity of Sm antibodies to various sDMA-containing epitopes, combined with a dominant antigenic specificity, provides mechanistic insight in the potential link between viral infection and tolerance breach in anti-Sm+ SLE patients.

## INTRODUCTION

Systemic lupus erythematosus (SLE) is a chronic systemic autoimmune disease characterized by immune cell dysfunction, leading to widespread inflammation and tissue damage [1]. Autoreactive B cells are central to SLE pathophysiology, a concept further emphasized by the recent successes of anti-CD19 CAR-T cell therapy in achieving clinical remission [2]. The antibodies produced by these self-reactive B cells are known as anti-nuclear antibodies (ANAs), a heterogeneous group of autoantibodies that recognize different chromatin components (e.g. double stranded DNA (dsDNA), histones) or RNA-associated proteins (e.g. Smith (Sm), Ro60). Although ANAs can be present in various autoimmune diseases, anti-dsDNA and anti-Sm antibodies are most specific to SLE, and have therefore been included among the EULAR-American College of Rheumatology (ACR) criteria for classification of this disease [1,3].

The Sm protein is composed of eight subunits with molecular weights varying between 9 and 29.5 kDa. It is part of the spliceosome complex, involved in the processing of nuclear pre-mRNA [4]. Using peptide screens, several antigenic epitopes have been characterized on the Sm protein, most prominently within the C-terminal domains of SmB/B’, SmD1, and SmD3 [5–8]. Intriguingly, these regions are rich in RGG/RG repeats, which mediate RNA binding, and contain arginines (Arg) that can undergo post-translational modification (PTM) to form symmetrical dimethylarginine (sDMA) residues *in vivo* [9,10]. These modifications appear to have immunological relevance, as subsequent research showed strong recognition of SmD1_95-119_ and SmD3_108-122_ peptides, each containing sDMA residues [11,12]. Importantly, both studies demonstrated that the introduction of an sDMA residue was essential for recognition by anti-Sm autoantibodies. Nonetheless, the context of surrounding amino acids to which anti-Sm autoantibodies binds remains uncertain, as their recognition has thus far been evaluated only in the context of the Sm protein complex.

By comparison, autoantibodies to PTMs, particularly citrullinated, carbamylated, and acetylated epitopes, form a major component of the autoimmune response in rheumatoid arthritis (RA) and are strongly associated with disease development [13]. These PTM-directed autoantibodies, also referred to as anti-modified protein antibodies (AMPA), are often described as “promiscuous”, reflecting their ability to cross-react with multiple citrullinated epitopes [14]. Whether anti-Sm autoantibodies in SLE share this feature and recognize sDMA residues across different epitopes has not been established. To address this gap, we sought to clarify the specificity and cross-reactivity of anti-Sm, focusing on the role of sDMA residues. In the current study, we show that antibodies targeting sDMA are common in anti-Sm+ SLE patients and represent the vast majority of the anti-Sm autoantibody repertoire. We further demonstrate that anti-sDMA IgGs are highly cross-reactive and can also bind to sDMA on multiple other targets, including several virus-derived epitopes, thus representing a prototypical anti-PTM response.

## MATERIALS AND METHODS

### Patients and healthy subjects

For the primary cohort, serum was obtained from 24 patients (12 anti-SmD+ and 12 anti-SmD-) with a clinical diagnosis of SLE, and all but one fulfilling the 2019 EULAR/ACR classification criteria for SLE, from the Rheumatology outpatient clinic at the Leiden University Medical Centre (LUMC; Leiden, The Netherlands) [3]. The second cohort comprised 40 additional SLE patients (20 anti-SmD+ and 20 anti-SmD-), all meeting the 2019 EULAR/ACR classification criteria for SLE, obtained from the neuropsychiatric (NP-) SLE biobank at the LUMC [3,15]. It is important to note that only approximately one-third of the SLE patients in this cohort were diagnosed with NP-SLE. Information regarding patient characteristics can be found in Supplementary Table 1 and 2. No information on ethnicity was available in the primary cohort. Anti-SmD positivity was established through clinical diagnostic fluorescence enzyme immunoassay (FEIA) testing and subsequently confirmed by in-house Sm protein enzyme-linked immunosorbent assays (ELISA), as detailed below. Clinical and routine laboratory data were collected within one week of serum collection for both cohorts. None of the SLE patients in the primary cohort had been treated with belimumab or rituximab in the 12 months preceding serum collection. The study with the primary cohort was approved by the Leiden Den Haag Delft medical ethics committee, while the study with the secondary cohort was approved by the LUMC biobank committee. Serum from 32 healthy donors (HDs) was obtained from the LUMC Voluntary Donor Service, with approval from the LUMC Biobank committee. All subjects have given written informed consent.

### Peptide synthesis and integrity identification

Biotinylated peptides were synthesized by the Peptide and Tetramer Facility Immunology at the LUMC as previously described [16] (detailed sequences in Supplementary Table 3). Briefly, peptides were synthesized using standard solid-phase 9-fluorenylmethoxycarbonyl (Fmoc) chemistry on an automated peptide synthesizer. After chain assembly, peptides were cleaved from the resin with trifluoroacetic acid (TFA), purified by reverse-phase high-performance liquid chromatography (HPLC) and lyophilized. Peptide cyclization was performed via thioether bond formation in 20 mM Tris-HCl buffer (pH ∼8.0). Lyophilized peptides were added slowly to 10 mL of buffer and incubated at RT for 3 hours, and cyclization was monitored by analytical reverse-phase ultra-performance liquid chromatography (UPLC). The mixture was acidified with 90% HOAc and purified on a Biotage Selekt system using a Sfär C18 Duo 100 Å, 30µm, 6g column (Biotage, FSUD-0401-0006). The integrity of the synthesized peptides after purification was assessed by UPLC using an Acquity system (Waters), and the exact masses were determined by mass spectrometry (MS) on a Microflex instrument (Bruker) and compared with the calculated values.

### Enzyme-linked immunosorbent assays

Standard capacity 384-well streptavidin microplates (Microcoat, 604500) were coated with 1 μg/mL of the respective peptide in phosphate-buffered saline (PBS; pH∼7.6) with 0.1% bovine serum albumin (BSA) for 1 hour at 37°C. Plates were washed and incubated with sera (diluted 1:200) in PBTT (PBS/1%BSA/50mM Tris/0.05% Tween-20, pH∼8.0), for 1 hour at 37°C. All washing steps were performed by washing 4x with 0.05% Tween-20 (Sigma-Aldrich, P1379) in PBS. After incubation, the plates were washed and incubated with 1 μg/mL anti-IgG horse-radish peroxidase (Bethyl; A80-104P).

IgGs against specific ANAs were determined using the protocol described above, with some adjustments. Clear flat bottom 384-well microplates (Corning, 3640) were coated overnight at 4°C with either 0.8 µg/mL bovine non-recombinant Sm (Diarect, A175), 1 µg/mL salmon sperm dsDNA (Invitrogen, 15632011), 0.5 µg/mL bovine non-recombinant histone (Diarect, A311), 0.5 µg/mL recombinant U1-snRNP 68/70 kDa (Diarect, A130) or 0.5 µg/mL recombinant Ro60/SS-A (Diarect, A174). Prior to serum incubation, plates were blocked with PBT (PBS/1%BSA/50mM Tris, pH∼8.0) for 1 hour. All procedures were conducted at room temperature (RT). In-house ELISAs for specific ANAs were previously developed and validated [17].

All ELISAs were developed using ABTS (Sigma-Aldrich, A1888) and H_2_O_2_ (Merck, 1072100250) and measured on an iMark Microplate Absorbance Reader (BioRad). Samples were assessed with technical duplicates.

### ELISA-based inhibition assays

Inhibition assays were performed as described previously, with some modifications [18]. Briefly, sera (diluted 1:200) were preincubated with indicated molar concentrations of blocking peptide for 2 hours at 37°C, prior to addition to the ELISA plates as described above. Inhibition percentages were calculated by subtracting the mean background OD (from HDs) and dividing the OD at the highest blocking concentration by the OD of the unblocked sample, with negative inhibition values were set to 0%.

### Antibody affinity purification

NeutrAvidin Agarose resin (Thermo Scientific, 53151) was coupled with 0.1 µg/µL SmD3_108-122_ (sDMA or Arg) for 1 hour at RT with shaking. The resin was transferred to a 96-well filter plate (Orochem, OF1100) and incubated with serum (diluted 1:4) in PBS for 2 hours at RT with shaking. The unbound fraction was removed by centrifugation at 500×g for 1 minute. The bound fraction was eluted twice with 25 µL 100mM formic acid for 5 minutes with shaking at RT, followed by centrifugation at 500xg for 2 minutes. The eluate was neutralized to pH∼7 using Tris-HCL (pH∼9.2)

### Cell lines

EBV-infected JY or DR7 B cell lines (ATCC) or EBV-negative BCL-6/Bcl-xL transduced immortalized B cells (IMM) [19] were cultured in IMDM containing 10% FCS, 100U/mL penicillin/streptomycin, and 2mM GlutaMax. Whole cell extracts were made by incubation for 30 min on ice in PBS with 0.5% IGEPAL (Sigma-Aldrich, I3021), 150mM NaCl, and protease inhibitor cocktail (Roche, 11836170001). Lysates were centrifuged at 12.000×g for 20 minutes at 4°C, supernatants were used for further analysis.

### Western blot

A total of 750 μg cell lysate was diluted in 4x Laemmli buffer (Bio-Rad, 1610747) and heated for 5 minutes at 95°C. Samples were run alongside a PageRuler Plus Prestained Protein Ladder (Thermo Fisher, 26619) on a 4–15% SDS–polyacrylamide gel (Bio-Rad, 4561084) and subsequently transferred to a PVDF membrane. The blots were blocked overnight in 5% milk/PBS/0.05% Tween at 4°C and subsequently incubated for 1 hour with either an anti-sDMA antibody (diluted 1:500; Sigma-Aldrich, 07-413; clone SYM11), anti-EBNA-1 antibody (diluted 1:1.000; Sigma-Aldrich, MABF2800; clone 1H4), or affinity purified anti-sDMA IgGs (∼1 μg/mL). After three washes with PBS/0.05%Tween, blots were incubated with either anti-rat IgG-AF680 (diluted 1:5.000; Invitrogen, A-21096), HRP-conjugated anti-rabbit Ig (diluted 1:1.000; DAKO, P0448) or HRP-conjugated anti-human IgG (diluted 1:1.000; DAKO, P0214). Blots were visualized using Pierce ECL substrate according to manufacturer’s instructions. Images were taken using iBright FL1500 Imaging System (Invitrogen).

### Statistical analysis

Statistical tests were performed in GraphPad Prism version 10.1.0. Paired comparisons were calculated using Wilcoxon signed-rank test and comparisons across multiple groups using a Friedman test, corrected for multiple testing using Bonferroni-Dunn method. *P* values <0.05 were considered statistically significant.

### Patient and public involvement

Patient feedback was obtained on the study design. Patients and/or the public were not involved in the conduct, reporting, or dissemination plans of this research.

## RESULTS

### sDMA is an essential constituent of the antigen recognized by anti-Sm antibodies irrespective of the flanking regions

To validate the requirement of an sDMA residue for recognition of the SmD3_108−122_ epitope by anti-Sm+ SLE patients [12], we assessed IgG reactivity against the full-length Sm protein as well as SmD3_108-122_ variants containing either sDMA, asymmetrical dimethyl arginine (aDMA), or unmodified arginine. We tested sera from anti-Sm+ SLE, anti-Sm− SLE, and HDs. Consistent with previous studies, the SmD3_108−122_ epitope was recognized exclusively by anti-Sm+ SLE patients, and only when containing sDMA (Figure 1A) [11,12]. One sDMA residue per peptide was sufficient for antibody binding as increasing the number of sDMA residues did not significantly enhance SmD3_108−122_ recognition by these patients (Supplementary figure 1). A scrambled peptide of the same amino acid composition was also included (sequence in Supplementary table 3) and surprisingly, was still specifically recognized by anti-Sm+ SLE sera (Figure 1B). These observations are intriguing as they point to anti-Sm autoantibodies directly targeting the PTM, similar to autoantibodies in RA directly recognizing citrulline (another PTM of arginine) in a variety of peptides [18,20]. Consequently, we continued to investigate whether anti-Sm antibodies represent a bona fide anti-PTM response, by generating a cyclic sDMA-containing peptide (CP-sDMA) modelled after the CCP4 peptide used in RA (substituting citrulline with sDMA) [21]. This peptide allows assessment of sDMA reactivity entirely independent of the context of the Sm protein. We tested sera from anti-Sm+ SLE, anti-Sm− SLE, and HDs and observed that the CP-sDMA was recognized exclusively by anti-Sm+ SLE patients (Figure 1C). These results suggest that anti-Sm+ patients contain IgG antibodies targeting sDMA in different contexts. To validate our findings, we assessed SmD3₁₀₈-₁₂₂ and CP-sDMA and reactivity in an independent second cohort (Figure 1D, E). Again, the CP-sDMA was recognized exclusively by anti-Sm+ SLE patients and to a similar extent as SmD3₁₀₈-₁₂₂, confirming that anti-PTM antibodies targeting sDMA are present in all anti-Sm+ patients from two cohorts.

**Figure 1:**
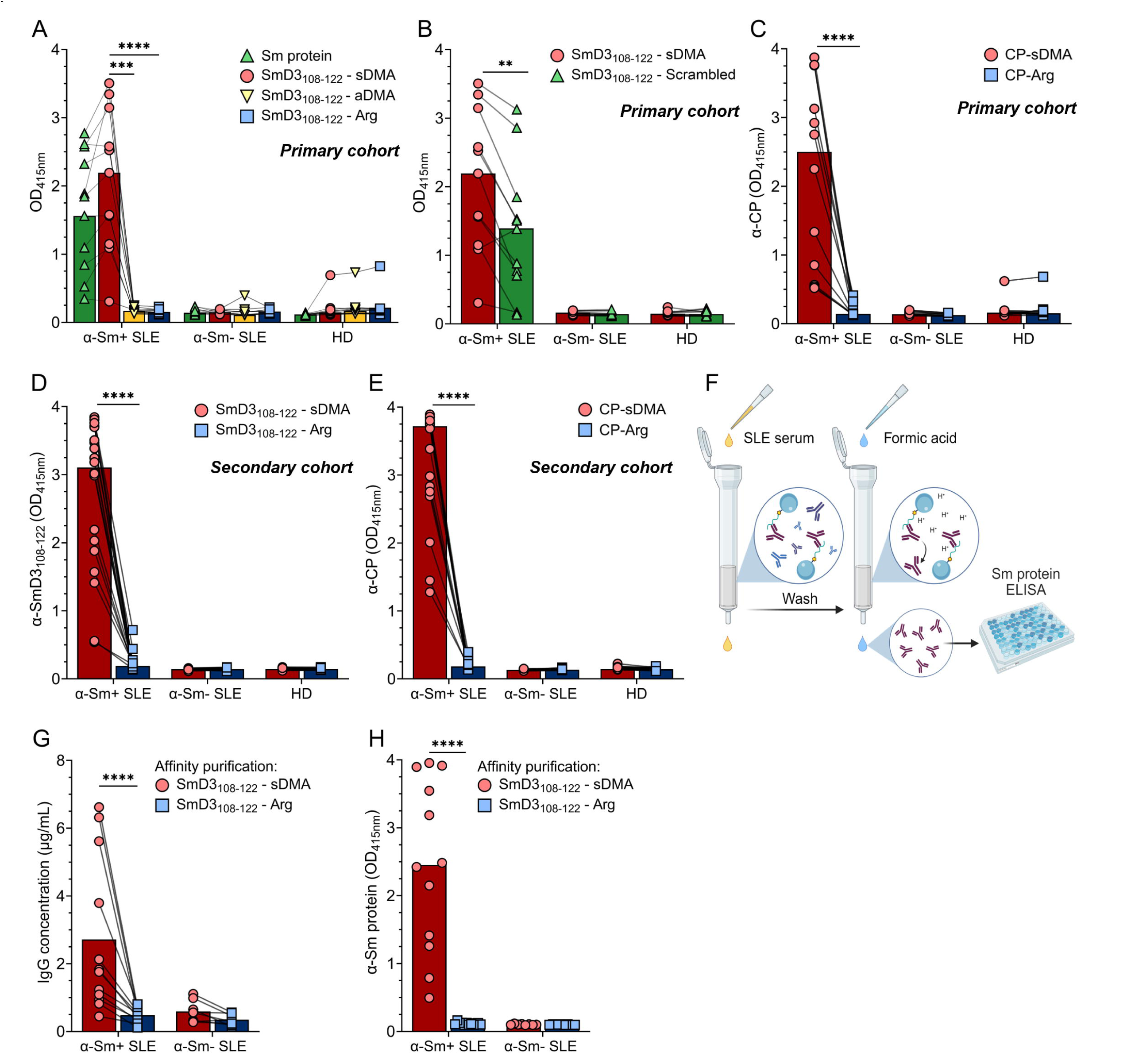
Characterization of anti-SmD3_108−122_ and other sDMA-containing peptides in anti-Sm+ SLE patients. Sera from anti-Sm+ (n=12), anti-Sm− (n=12) SLE patients, as well as HDs (n=12), were analysed by ELISA for IgG reactivity against: (A) the Sm protein and SmD3_108-122_ peptide variants containing either sDMA, aDMA, or unmodified arginine; (B) SmD3_108-122_ variant with a scrambled sequence; and (C) a cyclic peptide modified either with sDMA or an unmodified arginine. (D) Reactivity to SmD3_108-122_ and (E) a cyclic peptide, each tested in sDMA-modified or unmodified (arginine) form, was assessed in an independent secondary cohort of anti-Sm+ (n = 20) and anti-Sm− (n = 20) SLE patients and HDs (n = 20). (F) Schematic overview of antibody affinity purification, where SmD3_108-122_ binding antibodies were isolated from anti-Sm+ (n=12) and anti-Sm− (n=8) SLE sera using either sDMA or arginine control variants. (G) IgG concentration after affinity purification and (H) antibody binding to the Sm protein after affinity purification were measured by ELISA. All bar graphs show the median with lines connecting data points from individual samples. One representative experiment of three is shown. Data were analysed using simple linear regression or a Friedman test; ** P≤ 0.01, *** P ≤ 0.001, **** P ≤ 0.0001. Sm, Smith; SLE, Systemic Lupus Erythematosus; sDMA, symmetrical dimethylarginine; aDMA, asymmetrical dimethylarginine; Arg, arginine; HD, healthy donor; OD_415_, optical density at 415 nm; CP, cyclic peptide.

Next, to evaluate whether IgGs targeting sDMA-containing peptides recognize the full-length Sm protein, we isolated antibodies by affinity purification with SmD3_108-122_ (sDMA or Arg) (Figure 1F). Significantly higher amounts of IgGs were isolated from anti-Sm+ SLE patients using the sDMA-modified SmD3_108-122_ peptide compared to the unmodified control (Figure 1G). Only the anti-sDMA antibodies from anti-Sm+ SLE patients exhibited binding to the full-length Sm protein (Figure 1H). Overall, our findings indicate that IgG from anti-Sm+ SLE patient sera recognize the SmD3_108-122_ epitope through its sDMA residue, and that these anti-sDMA IgGs are capable of targeting the full-length Sm protein. Importantly, sDMA recognition can occur in the context of sequences independent from Sm.

### sDMA forms the predominant target of the anti-Sm response in SLE patients

We noticed a strong correlation between the levels of IgG reactive to SmD3_108−122_ and full-length Sm (Figure 2A). Therefore, we sought to determine to what extent IgGs targeting sDMA-containing epitopes contribute to the overall anti-Sm IgG response in SLE patients. To assess this, we conducted an inhibition assay incubating anti-Sm+ SLE serum with various concentrations of SmD3_108-122_ (sDMA or Arg) prior to detection of anti-Sm full-length protein by ELISA (Figure 2B). In order to confirm that anti-sDMA IgGs were effectively blocked, we first tested the blocked serum for recognition of SmD3_108-122_–sDMA. Blocking with SmD3_108-122_–sDMA effectively inhibited recognition in a dose-dependent manner in most samples, whereas blocking with SmD3_108-122_–Arg had no effect on recognition. At the highest concentration of blocking peptide, nearly all sDMA-specific IgGs were successfully blocked (Supplementary figure 2). Blocking anti-SmD3_108-122_–sDMA IgG resulted in a significant reduction in Sm protein recognition (Figure 2C), with an average inhibition of ∼90% (Figure 2E). No effect in Sm recognition was observed when anti-Sm+ sera were blocked with the arginine control peptide (Figure 2D) or in anti-Sm− SLE patients and HDs (Supplementary Figure 3). Furthermore, to confirm that this inhibition of Sm recognition depends exclusively on the sDMA residue, we performed a similar inhibition assay using the CP-sDMA (or arginine control) (Figure 2F), resulting once more in an average inhibition of ∼90% (Figure 2G-I). Importantly, blocking sDMA did not affect the recognition of other ANAs (dsDNA, histone, RNP70 and Ro60/SS-A), confirming the specificity of sDMA antibodies for the anti-Sm response (Supplementary figure 4). For comparison, anti-Sm+ SLE sera were also incubated with the cross-reactive PPPGMRPP peptide, previously described as an early target in the anti-Sm response [5]. Notably, the PPPGMRPP peptide was not differently recognized by anti-Sm+ SLE sera compared to anti-Sm− SLE or HDs (Figure 2J) and inhibition with this peptide had no effect on Sm protein recognition (Figure 2K). Taken together, our findings show that the sDMA residue is an essential constituent of the antigens targeted by the majority of the anti-Sm repertoire.

**Figure 2:**
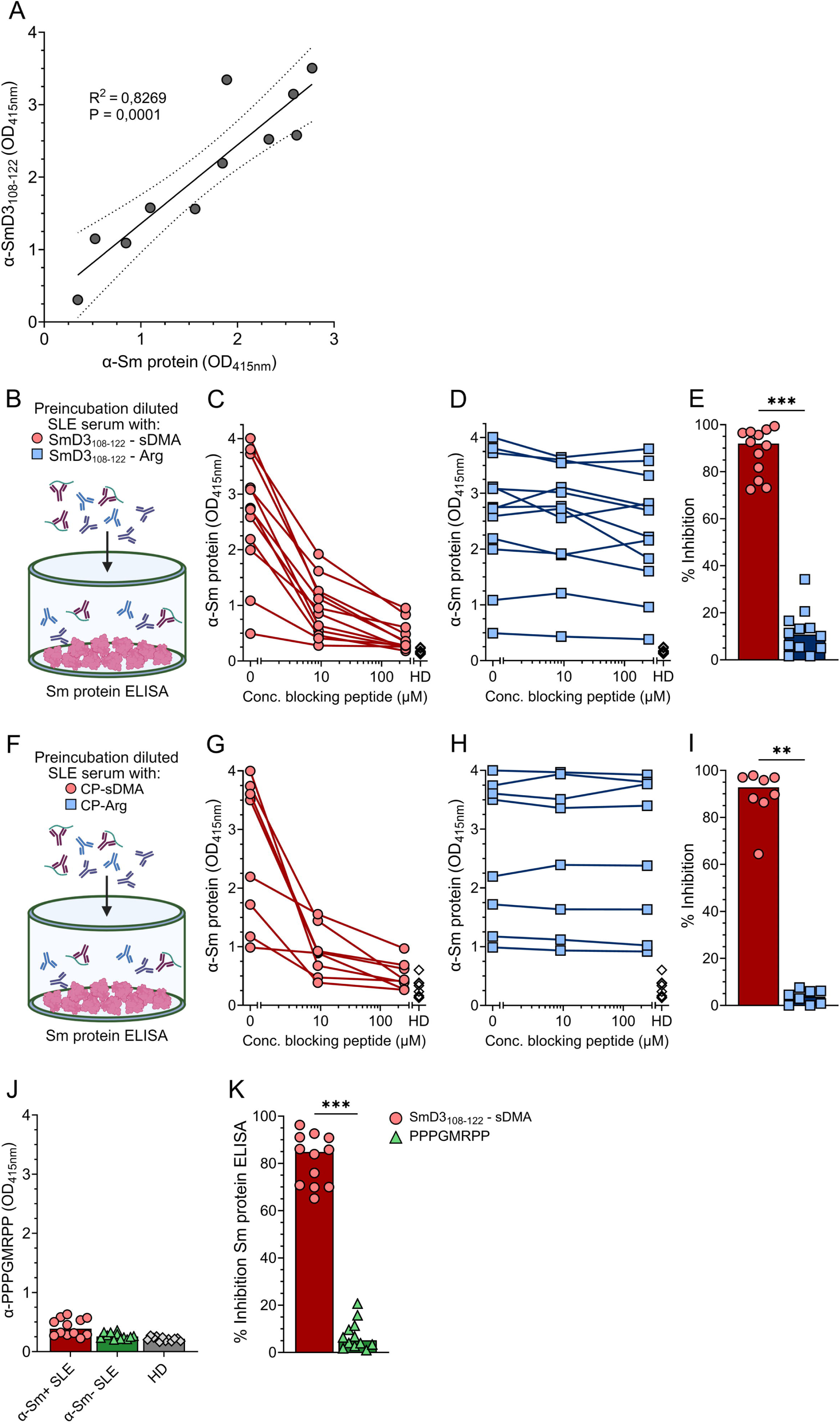
sDMA forms the predominant target of the anti-Sm response in SLE patients. (A) Correlation of anti-SmD3_108-122_-sDMA OD_415_ with anti-Sm OD_415_ in anti-Sm+ SLE patients (n=12) (data from Figure 1A). (B/F) Schematic overview of the inhibition assay, where sera of anti-Sm+ SLE patients were incubated with increasing concentrations of blocking (C-E) SmD3_108-122_ peptide (n=12) or (G-I) cyclic peptide (n=8) (sDMA (red circles) or arginine control (blue squares)) and subsequently measured on Sm protein ELISA. (J) ELISA showing PPPGMRPP peptide binding by sera from anti-Sm+ (n=12), anti-Sm− (n=12) SLE patients, and HDs (n=12). (K) Inhibition assay where sera of anti-Sm+ SLE patients (n=6) were incubated with increasing concentrations of PPPGMRPP or SmD3_108-122_-sDMA and subsequently measured on Sm ELISA. (E/I/J) Bar graphs show percent inhibition at 250 µM relative to the unblocked condition. All bar graphs show the median and line graphs connect individual measurements from the same patient sample. One representative experiment of three independent repetitions is shown. Data were analysed using simple linear regression or a Wilcoxon signed-rank test; * P ≤ 0.05, ** P ≤ 0.01, *** P ≤ 0.001. Sm, Smith; SLE, Systemic Lupus Erythematosus; sDMA, symmetrical dimethylarginine; Arg, arginine; OD_415_, optical density at 415 nm; ELISA, enzyme-linked immunosorbent assay.

### Anti-Sm IgG is cross-reactive to viral sDMA-containing epitopes

In addition to the Sm protein, emerging evidence suggests that several viral proteins carry sDMA residues as well [22–24]. To determine whether anti-sDMA IgG from anti-Sm+ SLE patients can recognize viral sDMA-containing epitopes, we synthesized peptides of several viral proteins, each containing sDMA at a specified position or unmodified arginine as control. We assessed peptides derived from Epstein–Barr virus (EBV) nuclear antigen 1 (EBNA1) [25], EBNA2 [26,27], Hepatitis C virus (HCV) non-structural protein 3 (NS3) [28], and infectious bursal disease virus (IBDV) RNA polymerase VP1 [29]. All sDMA-containing viral peptides were specifically recognized by anti-Sm+ SLE patients (Figure 3A-D). Similar to the SmD3_108-122_ epitope, recognition was dependent on the presence of sDMA. We also occasionally observed recognition of viral peptides in anti-Sm− SLE patients and HDs, but this reactivity was not sDMA-dependent, as the unmodified control was recognized equally well. Binding to the VP1_420-439_-sDMA peptide was modest compared to the other peptides (Figure 3D).

**Figure 3:**
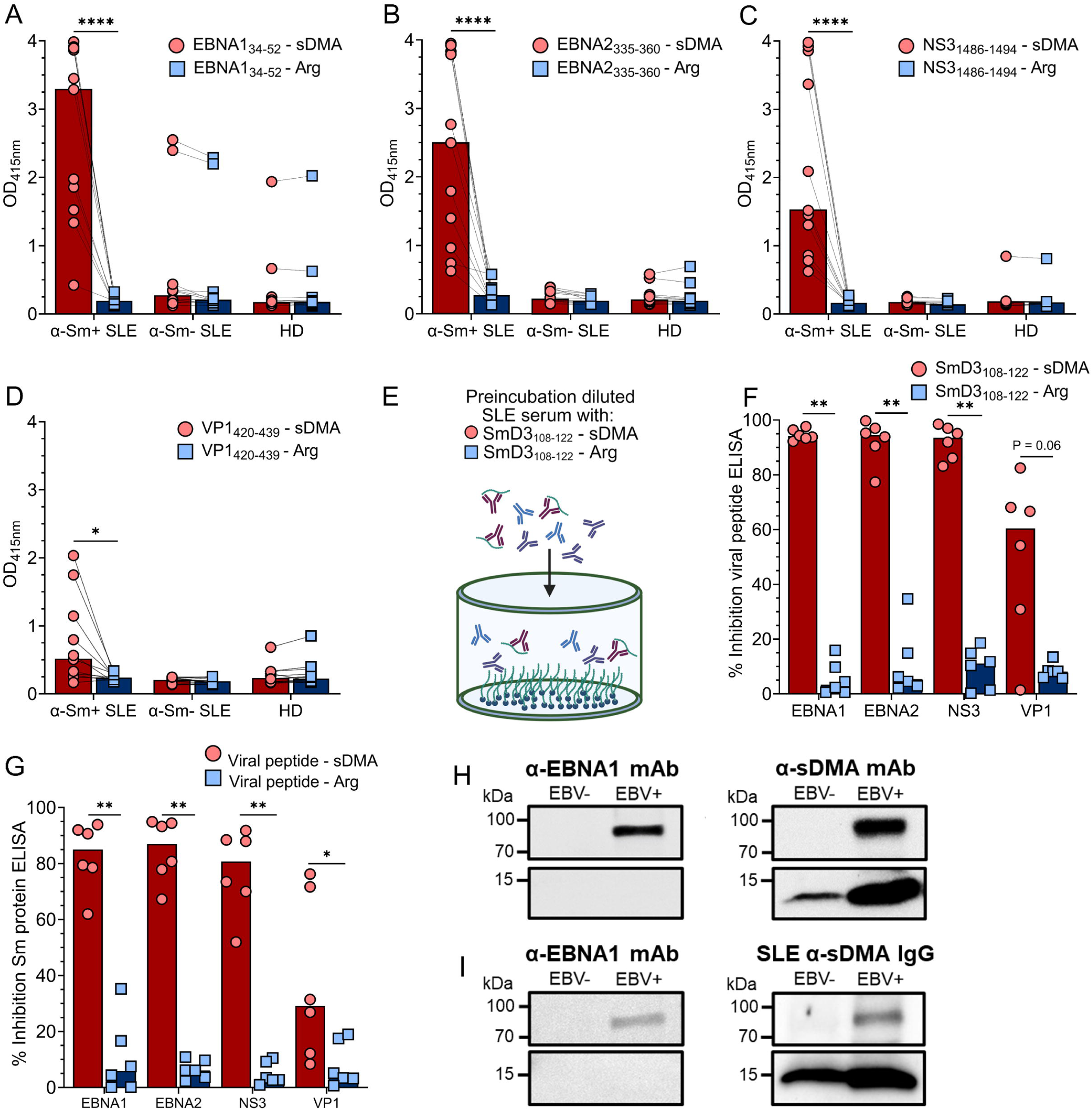
Anti-sDMA IgG antibodies from SLE patients are cross-reactive to multiple viral sDMA-containing epitopes. Sera from anti-Sm+ (n=12), anti-Sm− (n=12) SLE patients, as well as HDs (n=12), were analysed by ELISA for IgG reactivity against viral sDMA-containing epitopes or corresponding arginine controls: EBV-derived (A) EBNA1_34-52_ or (B) EBNA2_335-360_, (C) HCV-derived NS3_1486-1494_, and (D) IBDV-derived VP1_420-439_. (E) Schematic overview of an inhibition assay, where sera of anti-Sm+ SLE patients (n=6) were incubated with increasing concentrations of blocking SmD3_108-122_ peptide (sDMA or arginine control) and (F) subsequently measured on the respective sDMA-containing viral peptide ELISA. (G) Inhibition assay where sera of anti-Sm+ SLE patients (n=6) were incubated with respective viral peptides (sDMA or arginine control) and subsequently measured on Sm protein ELISA. (H) Lysates from EBV− IMM and EBV+ JY B cell lines were subjected to western blot analysis to detect EBNA1 and other sDMA-modified proteins, and (H) subsequently tested for recognition by sDMA-specific IgG antibodies purified from pooled anti-Sm+ SLE sera. (F/G) Bar graphs show percent inhibition at 250 µM relative to the unblocked condition. All bar graphs represent the median, with lines connecting paired samples. One representative experiment of three is shown. Data were analysed using a Wilcoxon signed-rank test; * P≤ 0.05, ** P ≤ 0.01, **** P ≤ 0.0001. Sm, Smith; SLE, Systemic Lupus Erythematosus; sDMA, symmetrical dimethylarginine; Arg, arginine; HD, healthy donor; OD_415_, optical density at 415 nm; EBV, Epstein-Barr virus; EBNA1, Epstein–Barr nuclear antigen 1; EBNA2, Epstein–Barr nuclear antigen 2; HCV, Hepatitis C virus; NS3, Non-structural protein 3; IBDV, Infectious Bursal Disease Virus; VP1, Viral Protein 1; IMM, BCL-6/Bcl-xL transduced immortalized; kDa, kilodalton.

To assess whether anti-sDMA IgGs are cross-reactive between SmD3_108-122_ and viral sDMA-containing peptides, we performed a similar inhibition assay as described previously by preincubating anti-Sm+ SLE serum was with various concentrations of SmD3_108-122_ (sDMA or Arg) before assessing binding to viral peptides by ELISA (Figure 3E). This resulted in an average inhibition ranging from 65% to 95%, indicating that anti-sDMA IgGs are highly cross-reactive to sDMA-containing peptides of various origins (Figure 3F, Supplementary Figure 5A-H). Conversely, we tested whether sDMA-containing viral peptides could similarly inhibit Sm protein recognition. Anti-Sm+ SLE sera were preincubated with the respective viral peptides (sDMA or Arg), which resulted in approximately 80–90% inhibition, indicating strong cross-reactivity between viral sDMA-containing epitopes and the Sm protein (Figure 3G, Supplementary Figure 5I-P).

Next, we sought to determine whether anti-sDMA IgG could also recognize sDMA residues within endogenous viral proteins. To this end, we selected EBNA1 as a model antigen. To confirm the presence of sDMA on endogenous EBNA1, we performed western blotting on lysates from EBV− and EBV+ B cell lines using anti-sDMA (SYM11) and anti-EBNA1 (1H4) monoclonal antibodies (Figure 3H, Supplementary figure 6A/B). The alignment of both signals at the same molecular weight supports the presence of sDMA on EBNA1. Next, blots were stained with polyclonal anti-sDMA IgG affinity-purified from multiple SLE patients (Figure 1H, I). In addition to the presumable SmD band at approximately 15 kDa, IgG binding was observed at the same molecular weight as the anti-EBNA1 band in the EBV+ lysate (Figure 3I, Supplementary figure 6C/D), suggesting that patient-derived anti-sDMA IgGs can bind endogenous EBNA1. Overall, our results indicate that IgGs targeting sDMA are highly cross-reactive to other sDMA-containing epitopes, including several viral-derived proteins.

## DISCUSSION

This study sought to deepen the understanding of the fine specificity and cross-reactivity of the anti-Sm autoantibodies response in SLE. Several antigenic targets have been identified on the Sm protein, with SmD3_108–122_ being recognized as a highly specific epitope that is also currently used in clinical diagnostics. [11,12] Building on previous studies, our findings now demonstrate that the sDMA residue is a primary target, largely independent of the AA sequence in which it is embedded. This conclusion is supported by the sustained recognition of scrambled SmD3_108–122_ and numerous other sDMA-containing peptides outside the structural context of the Sm antigen by anti-Sm+ SLE patient sera, including several viral-derived epitopes. Thus, the anti-Sm response represents a prototypic anti-PTM response, resembling the ACPA response and other AMPA responses described in RA [20].

The fine specificity of the anti-Sm response has been studied before by performing overlapping peptide screens using sera collected before and after SLE diagnosis. The PPPGMRPP motif of SmB/B’ was identified as a primary target of the early anti-Sm response and was suggested to be an initiating factor, though several other peptides were also recognized [5,8,30]. Although a potential initiating role cannot be ruled out, we did not observe a significant contribution of antibodies targeting PPPGMRPP to the anti-Sm antibody repertoire in established SLE patients from both cohorts. Importantly, peptide screens are inherently limited by the absence of PTMs in the peptides being used, thereby failing to capture the full spectrum of the antibody response against endogenous proteins. It is important to note that older studies already suggested that only native purified Sm, or Sm produced in baculovirus-infected insect cells, and not bacterially expressed Sm, was recognized by anti-Sm+ SLE serum [11]. These observations can now be explained by our findings, as sDMA is the major determinant of Sm recognition and symmetrical dimethylation by bacteria has not been observed to date [31]. In addition, sole recognition of sDMA residues also elucidates why anti-Sm antibodies were shown to recognize cross-reactive epitopes on both the B⍰/B and D subunits of the Sm complex, despite their limited AA sequence homology [32].

Here, we demonstrate for the first time that the majority (>90%) of anti-Sm IgGs specifically target sDMA residues, highlighting a central role of this PTM in the development of the SLE-specific B cell response to Sm. Additional human proteins, such as RNP70 and histones, have been suggested to carry sDMA [24,33], which may explain the strong association between anti-RNP70 and anti-Sm antibody responses in SLE patients [34,35]. Still, blocking anti-sDMA IgGs did not significantly affect the recognition of other ANAs, including recombinant RNP70 and non-recombinant purified histones, in anti-Sm+ SLE patients. Furthermore, purified anti-sDMA antibodies and monoclonals had restricted recognition of proteins from cell lysates on Western blot. Therefore, the presence and recognition of sDMA residues on other self-proteins may be limited, although still plausible, as a proteomic study showed that anti-Sm+ sera from SLE patients differentially recognized arginine-methylated proteins (including cellular nucleic acid binding protein (CNBP)), with methylation-dependent immunoreactivity reduced when methylation was inhibited [36], underscoring that methylarginine modifications can influence anti-Sm binding to non-Sm antigens. Why the sDMA-containing peptides play such a significant role in the recognition of Sm remains to be determined, but similarities between viral antigens and Sm-derived epitopes may offer a plausible explanation.

Numerous viruses have been linked to SLE pathogenesis, with a particular focus on EBV [37]. Molecular mimicry between viral and self-antigens could provide a trigger in the initiation of autoimmunity, and our results suggest strong mimicry between various sDMA-containing viral proteins and Sm. While some studies focused on a motif within EBV EBNA-1 (AA #398-404), which showed mimicry to the PPPGMRPP motif of SmB/B’ [8,30,38], we could not confirm reactivity to this peptide, nor did blocking with this peptide significantly reduce binding to complete Sm protein. Our findings of sDMA recognition as a determining factor in cross-reactivity between Sm and viral proteins provides new insight into the potential triggers of anti-Sm autoimmunity. Although further studies are required to substantiate this possibility, many viral proteins, including those from common viruses (EBNA-1/EBNA-2 from EBV, PB2 from Influenza A), have been reported to contain sDMA [22,23,26], and thus, different viruses may have the capacity to trigger a cross-reactive response targeting sDMA. Expression of the entire EBNA-1 protein in mice through an expression vector, induced IgGs that bound to non-recombinant Sm [39]. Since EBNA-1 expressed in mouse cells could acquire sDMA residues, our results may explain the cross-reactivity that was induced. We hypothesize that the cross-reactive anti-Sm response in SLE patients could arise from B cells that recognize sDMA-containing epitopes but receive T cell help specific for a viral antigen. When cross-reactivity is present within the B cell compartment, these B cells can receive help from foreign-reactive non-autoreactive T cells when they internalize and present peptides from the cross-reactive viral protein, thereby leading to break in B cell tolerance. Exposure to the viral antigen would be essential for this initial tolerance breach, as sDMA-reactive B cells would otherwise not receive CD4+ T cell help. Notably, anti-Sm positivity in SLE is strongly associated with particular combinations of HLA class II haplotypes [40], underscoring the central role of CD4+ T cells in shaping these pathogenic B cell responses. Interestingly, antibodies against sDMA-containing SmD3_108–122_ peptide were not detected during active viral infection in otherwise healthy individuals, such as Cytomegalovirus, HCV, or EBV [12]. This suggests that viral infection alone is not sufficient to trigger an anti-sDMA response, and that other factors in patients with SLE likely contribute to loss of tolerance against sDMA-containing self-antigens [37]. Recent work has demonstrated that SLE patients have higher numbers of EBV-infected B cells and that these cells are more activated [41]. Whether this is the cause or consequence of SLE remains to be determined. However, increased exposure to EBV-derived proteins containing sDMA may contribute to loss of tolerance and may thereby provide a potential bridge between EBV infection and the emergence of Sm autoreactivity in SLE.

Despite the evidence provided in this study, several limitations need to be considered. In this study, we primarily relied on synthetic peptides and included only one model viral protein antigen: EBNA1. Although we demonstrate cross-reactivity to various viruses at the peptide level, evidence at viral protein level remains indirect. It therefore remains speculative at this stage whether anti-sDMA IgG autoantibodies are truly cross-reactive to viral proteins *in vivo* and whether such cross-reactivity contributes to the initiation of the anti-Sm response in SLE.

In conclusion, our findings highlight the central role of sDMA residues in the development of anti-Sm autoantibodies in SLE, thereby representing a prototypic anti-PTM response. Furthermore, the high cross-reactivity of Sm antibodies to various sDMA-containing epitopes reveals a novel mechanistic perspective on the potential link between viral infection and tolerance breach in anti-Sm+ SLE patients.

## ACKNOWLEDGEMENTS

Some figure components were generated using BioRender (see licensed resources at https://BioRender.com/694mb1q and https://BioRender.com/e8wwcm5) We gratefully acknowledge J.W. Drijfhout for his helpful contributions and discussions. This work was supported by the Dutch Arthritis Society [grant number 22-1-402].

