## Supplemental Figures for "Symmetrical Dimethylarginine as the Central Antigenic Determinant of Anti-Smith Autoantibodies in Systemic Lupus Erythematosus"

| Patient demographics | α-Sm+ SLE (n=12) | α-Sm- SLE (n=12) | Healthy controls (n=12) |
|---|---|---|---|
| Age (yrs) | 43 (21-65) | 56 (27-82) | 36 (24-66) |
| Male / female (% female) | 2/10 (83%) | 1/11 (92%) | 2/10 (83%) |
| Disease duration (yrs) | 14 (1-32) | 19 (6-48) | N/A |
| Age at diagnosis (yrs) | 30 (13-51) | 35 (11-54) | N/A |
| **SLEDAI-2k** | 6 (2-20) | 9 (1-12) | N/A |
| **Clinical symptoms** | | | |
| Mucocutaneous (%) | 9/12 (75%) | 7/12 (58%) | N/A |
| Arthritis (%) | 2/12 (17%) | 6/12 (50%) | N/A |
| Renal (%) | 1/12 (8%) | 1/12 (8%) | N/A |
| Other (%) | 1/12 (8%) | 4/12 (33%) | N/A |
| **Haematology** | | | |
| Leukocytes, $\times 10^9$/L | 4.89 (3.54-8.63) | 6.24 (2.62-8.47) | N/A |
| Thrombocytes, $\times 10^9$/L | 248 (179-328) | 216 (86-426) | N/A |
| Lymphocytes, $\times 10^9$/L | 1.07 (0.21-2.17) | 0.78 (0.45-1.92) | N/A |
| C3, g/L | 0.79 (0.56-1.25) | 0.95 (0.76-1.14) | N/A |
| C4, mg/L | 150 (69-248) | 168 (36-459) | N/A |
| **Autoantibodies** | | | |
| Anti-dsDNA (%) | 9/12 (75%) | 4/12 (33%) | N/A |
| Anti-Sm (%) | 12/12 (100%) | 0/12 (0%) | N/A |
| IU/ml (Cut-off positivity: ≥10) | 72 (12-330) | 0.6 (0-3.5) | N/A |
| Anti-RNP70kD (%) | 1/12 (8%) | 1/12 (8%) | N/A |
| Anti-U1RNP (%) | 6/12 (50%) | 2/12 (17%) | N/A |
| Anti-SS-A (%) | 4/12 (33%) | 2/12 (17%) | N/A |
| Anti-SS-B (%) | 3/12 (25%) | 3/12 (25%) | N/A |
| **Current medication** | | | |
| PDN (%) | 7/12 (58%) | 3/12 (25%) | N/A |
| HCQ (%) | 11/12 (92%) | 7/12 (58%) | N/A |
| MMF (%) | 2/12 (17%) | 3/12 (25%) | N/A |
| Azathioprine (%) | 2/12 (17%) | 2/12 (17%) | N/A |
| MTX (%) | 1/12 (8%) | 1/12 (8%) | N/A |
| Other immunosuppressants[1] (%) | 1/12 (8%) | 1/12 (8%) | N/A |

**Supplementary Table 1: Baseline patient characteristics of the primary cohort.** The characteristics shown here are obtained during their initial assessment. All parameters are shown as the median and range or number and frequency within the patient group. None of the patients had used cyclophosphamide or rituximab/belimumab in the preceding 12 months. Sm, Smith; SLE, Systemic Lupus Erythematosus; PDN, prednisone; HCQ, hydroxychloroquine; MMF, mycophenolate mofetil; MTX, methotrexate.

[1] Other Immunosuppressants include Tacrolimus, NSAID and Ustekinumab.

| **Patient demographics** | **α-Sm+ SLE (n=20)** | **α-Sm- SLE (n=20)** | **Healthy controls (n=20)** |
|---|---|---|---|
| Age (yrs) | 37 (15-69) | 55 (33-73) | 45 (23-68) |
| Male / female (% female) | 2/18 (90%) | 3/17 (85%) | 2/18 (90%) |
| Disease duration (yrs) | 4 (0-21) | 4 (0-40) | N/A |
| Age at diagnosis (yrs) | 29 (14-69) | 43 (18-73) | N/A |
| **Ethnicity (self-identified)** | | | |
| Caucasian | 12/20 (60%) | 17/20 (85%) | N/A |
| Black / African descent | 5/20 (25%) | 2/20 (10%) | N/A |
| Asian | 2/20 (10%) | 0/20 (0%) | N/A |
| Mixed | 1/20 (5%) | 1/20 (5%) | N/A |
| **SLEDAI-2k** | 3 (0-15) | 2 (0-26) | N/A |
| **Clinical symptoms** | | | |
| Mucocutaneous (%) | 11/20 (55%) | 5/20 (25%) | N/A |
| Arthritis (%) | 2/20 (10%) | 1/20 (5%) | N/A |
| Renal (%) | 7/20 (35%) | 7/20 (35%) | N/A |
| Other (%) | 17/20 (85%) | 8/20 (40%) | N/A |
| **Haematology** | | | |
| Leukocytes, $\times 10^9$/L | 4.99 (2.01-17.0) | 8.99 (3.54-15.4) | N/A |
| Thrombocytes, $\times 10^9$/L | 221 (159-455) | 256 (131-608) | N/A |
| Lymphocytes, $\times 10^9$/L | 0.83 (0.57-1.15) | 0.98 (0.49-1.34) | N/A |
| C3, g/L | 0.85 (0.40-8.40) | 1.09 (0.60-1.40) | N/A |
| C4, mg/L | 111 (20-307) | 158 (15-328) | N/A |
| **Autoantibodies** | | | |
| Anti-dsDNA (%) | 10/20 (50%) | 3/20 (15%) | N/A |
| Anti-Sm (%) | 20/20 (100%) | 0/20 (0%) | N/A |
| Anti-RNP (%) | 13/20 (65%) | 1/20 (5%) | N/A |
| Anti-SS-A (%) | 12/20 (60%) | 5/20 (25%) | N/A |
| Anti-SS-B (%) | 2/20 (10%) | 1/20 (%) | N/A |
| **Current medication** | | | |
| PDN (%) | 12/20 (60%) | 13/20 (65%) | N/A |
| HCQ (%) | 15/20 (75%) | 11/20 (55%) | N/A |
| MMF (%) | 3/20 (15%) | 2/20 (10%) | N/A |
| Azathioprine (%) | 1/20 (5%) | 2/20 (10%) | N/A |
| MTX (%) | 1/20 (5%) | 1/20 (5%) | N/A |
| Other immunosuppressants[1] (%) | 5/20 (25%) | 3/20 (15%) | N/A |

**Supplementary Table 2: Baseline patient characteristics of the secondary cohort.**

The characteristics shown here are obtained during their initial assessment. All parameters are shown as the median and range or number and frequency within the patient group. IU/mL values for the anti-SmD FEIA were unavailable; only positive/negative status is reported. None of the patients had used cyclophosphamide or rituximab/belimumab in the preceding 12 months. Sm, Smith; SLE, Systemic Lupus Erythematosus; PDN, prednisone; HCQ, hydroxychloroquine; MMF, mycophenolate mofetil; MTX, methotrexate; FEIA, fluorescence enzyme immunoassay.

[1] Other Immunosuppressants include Tacrolimus, NSAID and Ustekinumab.

| Peptide name | Sequence |
|---|---|
| SmD3$_{108-122}$ – sDMA | UzAARG-sDMA-GRGMGRGNIF |
| SmD3$_{108-122}$ – Arg | UzAARGRGRGMGRGNIF |
| SmD3$_{108-122}$ – 2x sDMA (1) | UzAARG-sDMA-G-sDMA-GMGRGNIF |
| SmD3$_{108-122}$ – 2x sDMA (2) | UzAARG-sDMA-GRGMG-sDMA-GNIF |
| SmD3$_{108-122}$ – 3x sDMA | UzAARG-sDMA-G-sDMA-GMG-sDMA-GNIF |
| SmD3$_{108-122}$ – sDMA different Arg | UzAARGRGRGMG-sDMA-GNIF |
| SmD3$_{108-122}$ – Scrambled | UzGGM-sDMA-GGIRRAFGNAR |
| SmD3$_{108-122}$ – G113A | UzAARG-sDMA-ARGMGRGNIF |
| SmD3$_{108-122}$ – G111A, G113A | UzAARA-sDMA-ARGMGRGNIF |
| SmD3$_{108-122}$ – G113I | UzAARG-sDMA-IRGMGRGNIF |
| SmD3$_{108-122}$ – G111I, G113I | UzAARI-sDMA-IRGMGRGNIF |
| ENBA-1$_{34-52}$ – sDMA | UzRGGDNHGRGRG-sDMA-GRGRGGG |
| ENBA-1$_{34-52}$ – Arg | UzRGGDNHGRGRGRGRGRGGG |
| EBNA-2$_{335-360}$ – sDMA | UzQSSRGQSRGRG-sDMA-GRGRGKGKSRDKQ |
| EBNA-2$_{335-360}$ – Arg | UzQSSRGQSRGRGRGRGRGKGKSRDKQ |
| NS3$_{1486-1494}$ – sDMA | UzQRRG-sDMA-TGRG |
| NS3$_{1486-1494}$ – Arg | UzQRRGRTGRG |
| VP1$_{420-439}$ – sDMA | UzGEANCT-sDMA-QHMQAAMYILTR |
| VP1$_{420-439}$ – Arg | UzGEANCTRQHMQAAMYILTR |
| CP – sDMA (cyclic) | HQFRF-sDMA-GNleSRAACZO |
| CP – Arg (cyclic) | HQFRFRGNleSRAACZO |

**Supplementary Table 3: An overview of the peptide sequences.** U = Biotine, z = PEG3-spacer, Nle= norleucine, Z = 6-aminohexanoic acid, O = lys (biotine)-amide**.**

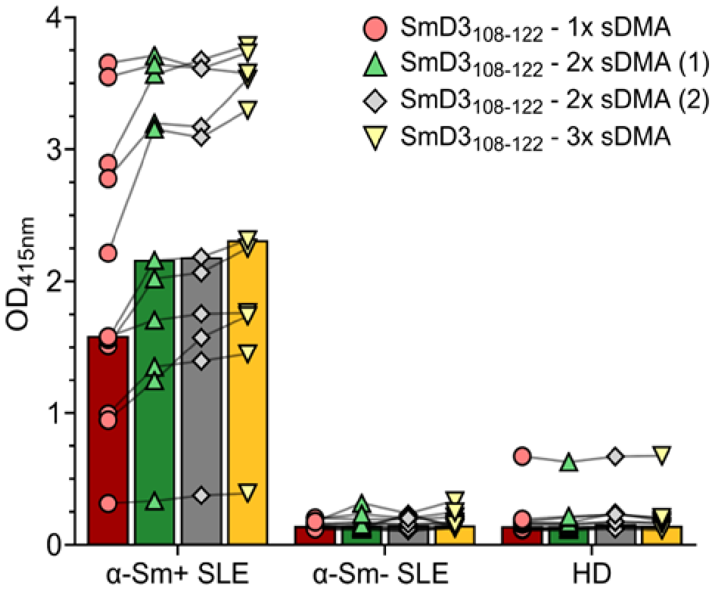


**Supplementary Figure 1: SmD3$_{108\text{-}122}$ epitope recognition containing multiple sDMA residues.**

Sera from anti-Sm+ (n=12), anti-Sm- (n=12) SLE patients, as well as HDs (n=12), were analysed by ELISA for IgG reactivity against SmD3$_{108\text{-}122}$ peptide variants containing one or multiple sDMA residues. All bar graphs represent the median, with lines connecting paired samples. One representative experiment of three is shown. Sm, Smith; SLE, Systemic Lupus Erythematosus; sDMA, symmetrical dimethylarginine; Arg, arginine; HD, healthy donor; $OD_{415}$, optical density at 415 nm; ELISA, enzyme-linked immunosorbent assay.

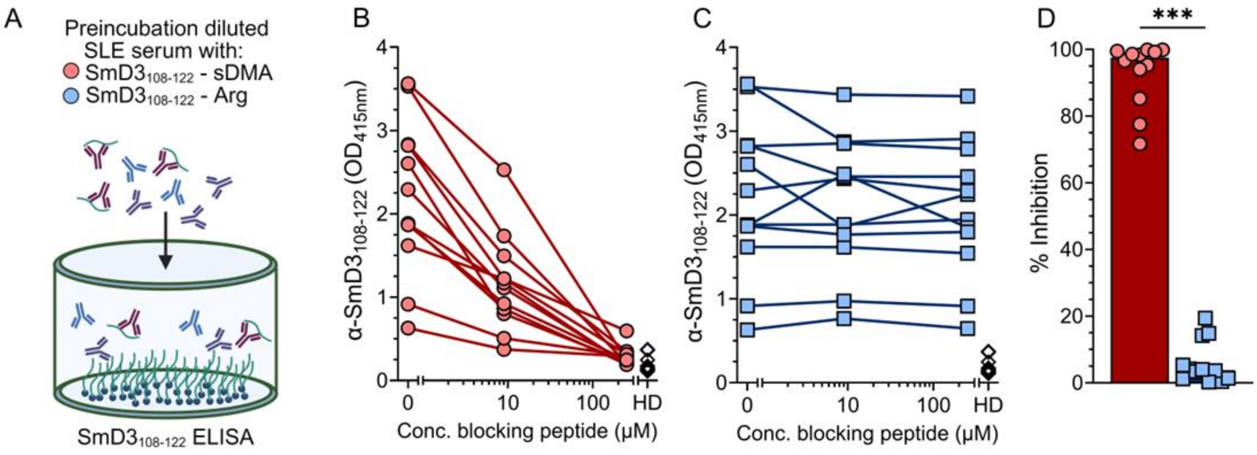


**Supplementary Figure 2: Validation of complete inhibition SmD3$_{108\text{-}122}$-sDMA directed IgG antibodies.** (A) Schematic overview of the inhibition assay, where sera of anti-Sm+ SLE patients were incubated with increasing concentrations of blocking (B) SmD3$_{108\text{-}122}$-sDMA or (C) SmD3$_{108\text{-}122}$-Arg peptide (n=12) and subsequently measured on SmD3$_{108\text{-}122}$-sDMA peptide ELISA. (D) Bar graph shows percent inhibition at 250 μM relative to the unblocked condition, indicating near 100% inhibition in most samples. One representative experiment of three independent repetitions is shown. Data were analysed using a Wilcoxon signed-rank test; *** P ≤ 0.001. Sm, Smith; SLE, Systemic Lupus Erythematosus; sDMA, symmetrical dimethylarginine; Arg, arginine; OD$_{415}$, optical density at 415 nm; HD, healthy donor; ELISA, enzyme-linked immunosorbent assay.

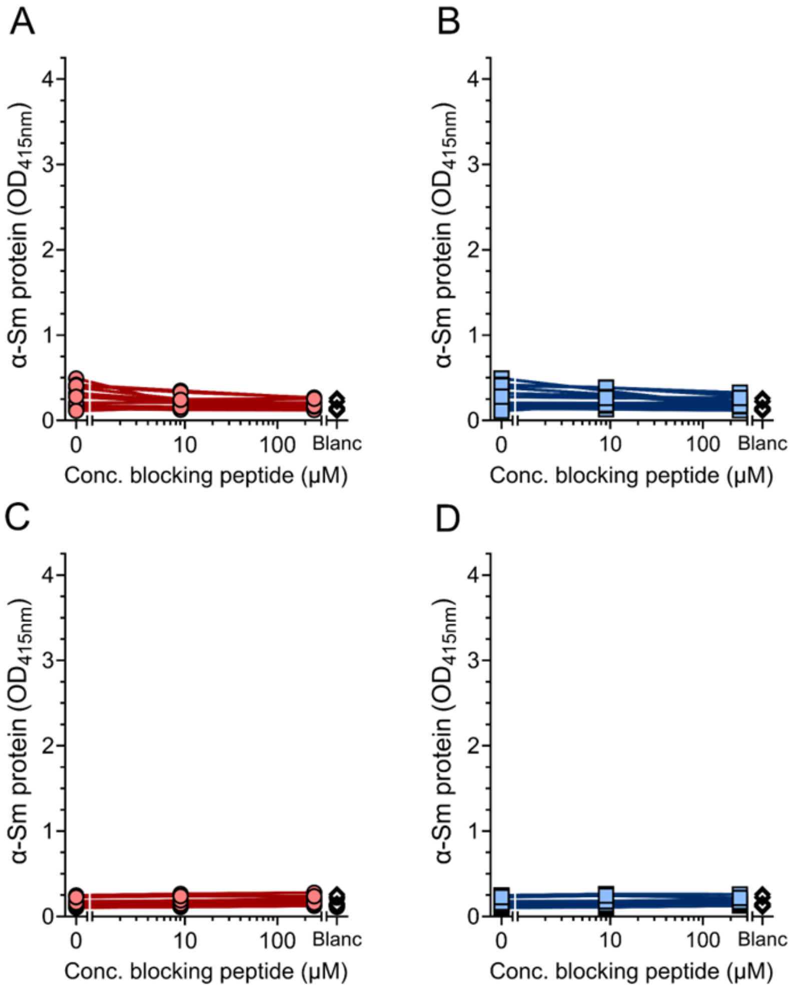


**Supplementary Figure 3: Inhibition assay in anti-Sm- SLE patients and HDs.** Sera from (A–B) anti-Sm- SLE patients (n=12) and (D–E) HDs (n=12) were incubated with increasing concentrations of blocking $SmD3_{108\text{-}122}$ peptide, containing either an sDMA (red circles) or unmodified arginine (blue squares), and tested on Sm protein ELISA. Line graphs connect individual measurements from the same patient sample. Sm, Smith; SLE, Systemic Lupus Erythematosus; sDMA, symmetrical dimethylarginine; $OD_{415}$, optical density at 415 nm; HD, healthy donor; ELISA, enzyme-linked immunosorbent assay.

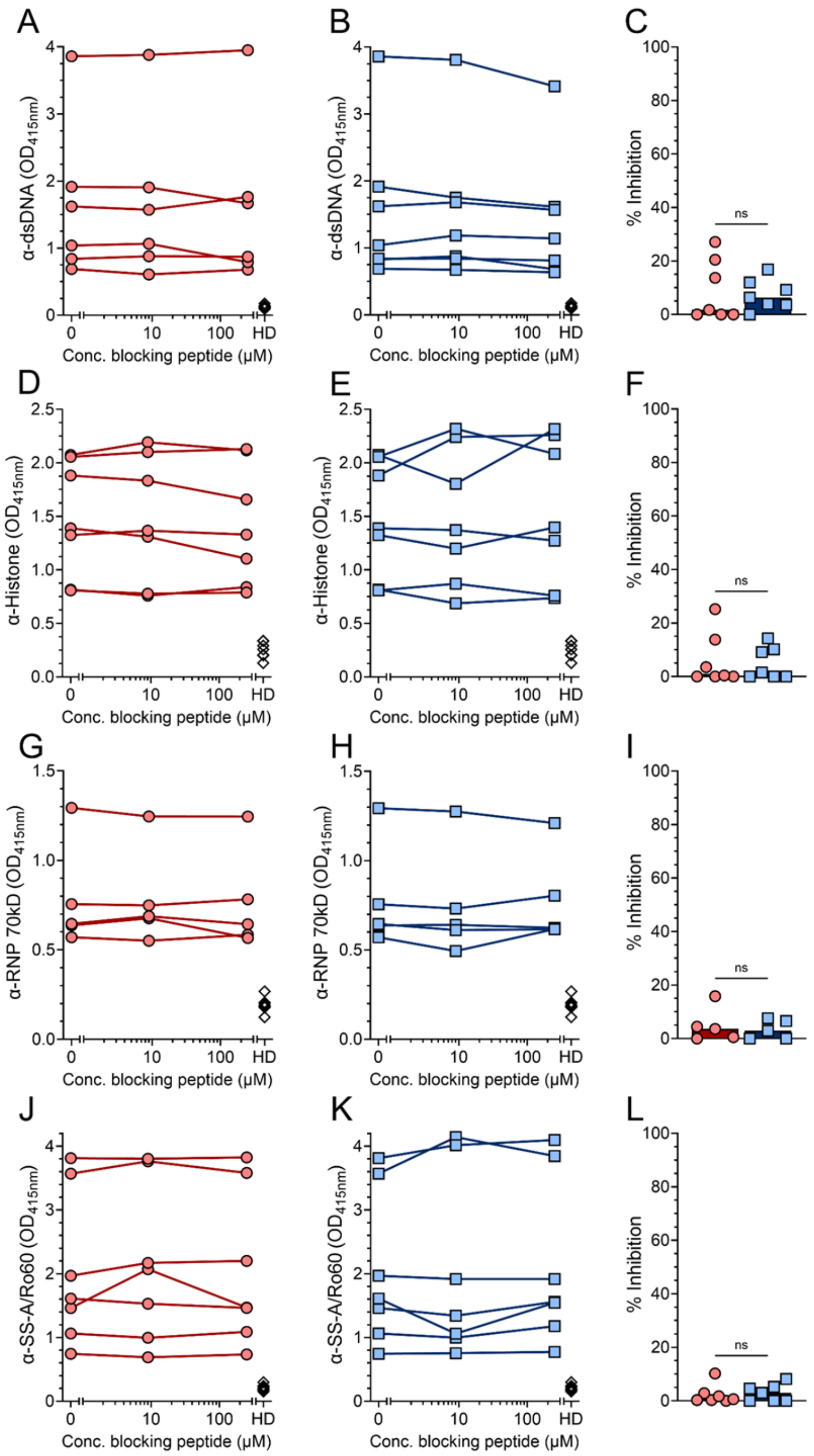


**Supplementary Figure 4: Effect of sDMA-IgG blocking on other ANA specificities in anti-Sm$^+$ SLE patients.** Sera from anti-Sm+ SLE patients (n=8) were incubated with increasing concentrations of blocking cyclic peptide, containing either an sDMA (red circles) or unmodified arginine (blue squares), and tested either (A-C) dsDNA, (D-F)

histone, (G-I) RNP70 or (J-L) SS-A/Ro60 ELISA. Only patients positive for IgG antibodies targeting the indicated nuclear antigen are shown. Line graphs connect individual measurements from the same patient sample. (C/F/I/L) Bar graphs show the median percent inhibition at 250 μM relative to the unblocked condition. Data were analysed using a Wilcoxon signed-rank test. Sm, Smith; dsDNA, double stranded DNA; SS-A, Sjogren's-syndrome-related antigen A; SLE, Systemic Lupus Erythematosus; sDMA, symmetrical dimethylarginine; $OD_{415}$, optical density at 415 nm; HD, healthy donor; ELISA, enzyme-linked immunosorbent assay; ns, not significant.

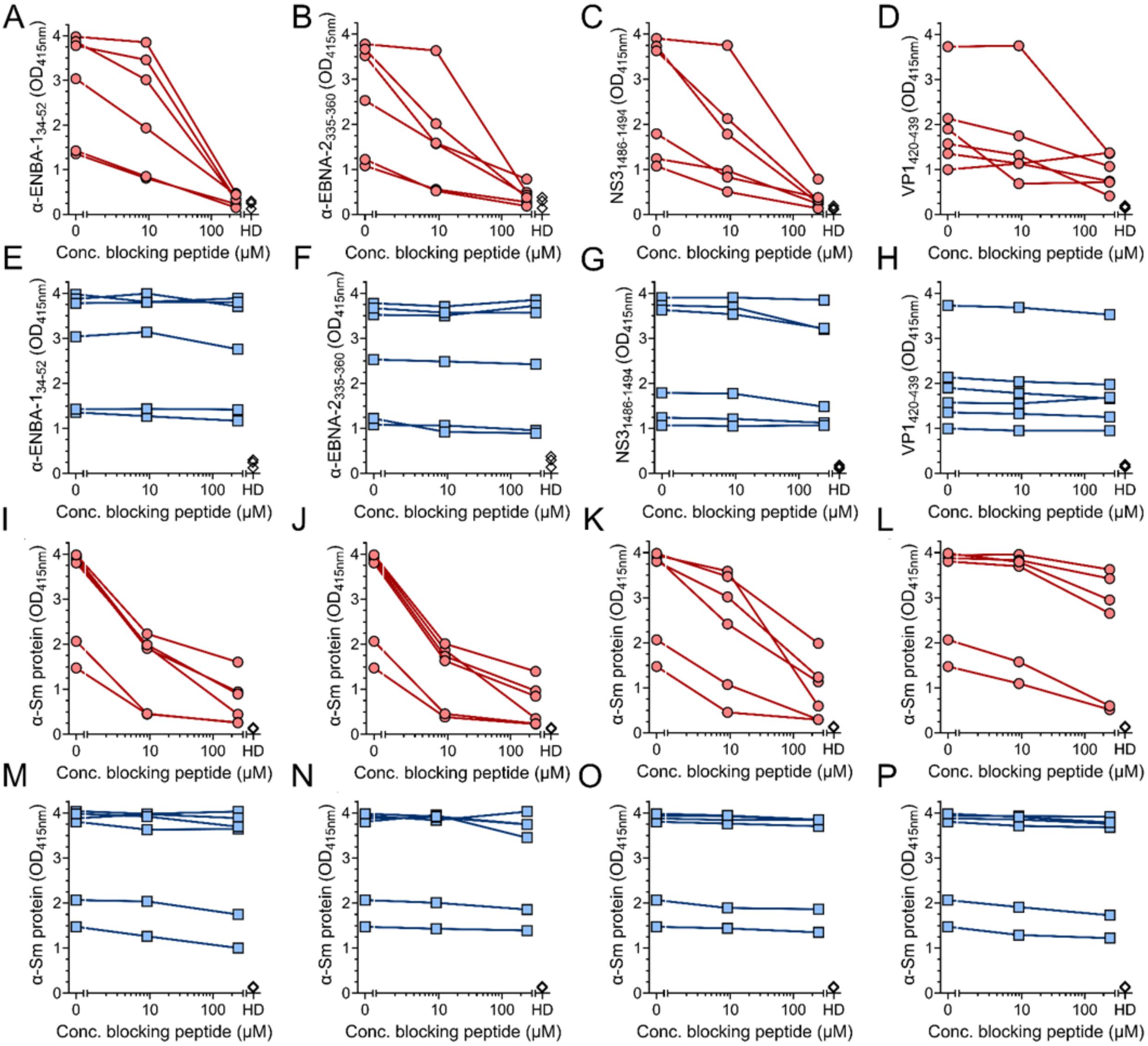


**Supplementary Figure 5: Line graphs of viral inhibition assay.** Line graphs corresponding to the bar graphs indicated in Figure 2F and 3J. Sera from anti-Sm+ SLE patients (n=6) were incubated with either increasing concentrations of (A-H) SmD3$_{108-122}$, (I/M) EBNA1$_{34-52}$, (J/N) EBNA2$_{335-360}$, (K/O) NS3$_{1486-1494}$, or (L/P) VP1$_{420-439}$ blocking peptide, containing either an sDMA (red) or unmodified arginine (blue) and tested on either the corresponding viral peptide or Sm protein ELISA. Line graphs connect individual measurements from the same patient sample. Sm, Smith; SLE, Systemic Lupus Erythematosus; sDMA, symmetrical dimethylarginine; HD, healthy donor; OD$_{415}$, optical density at 415 nm; ELISA, enzyme-linked immunosorbent assay; EBV, Epstein-Barr virus; EBNA1, Epstein–Barr nuclear antigen 1; EBNA2, Epstein–Barr nuclear antigen 2; HCV,

Hepatitis C virus; NS3, Non-structural protein 3; IBDV, Infectious Bursal Disease Virus; VP1, Viral Protein 1.

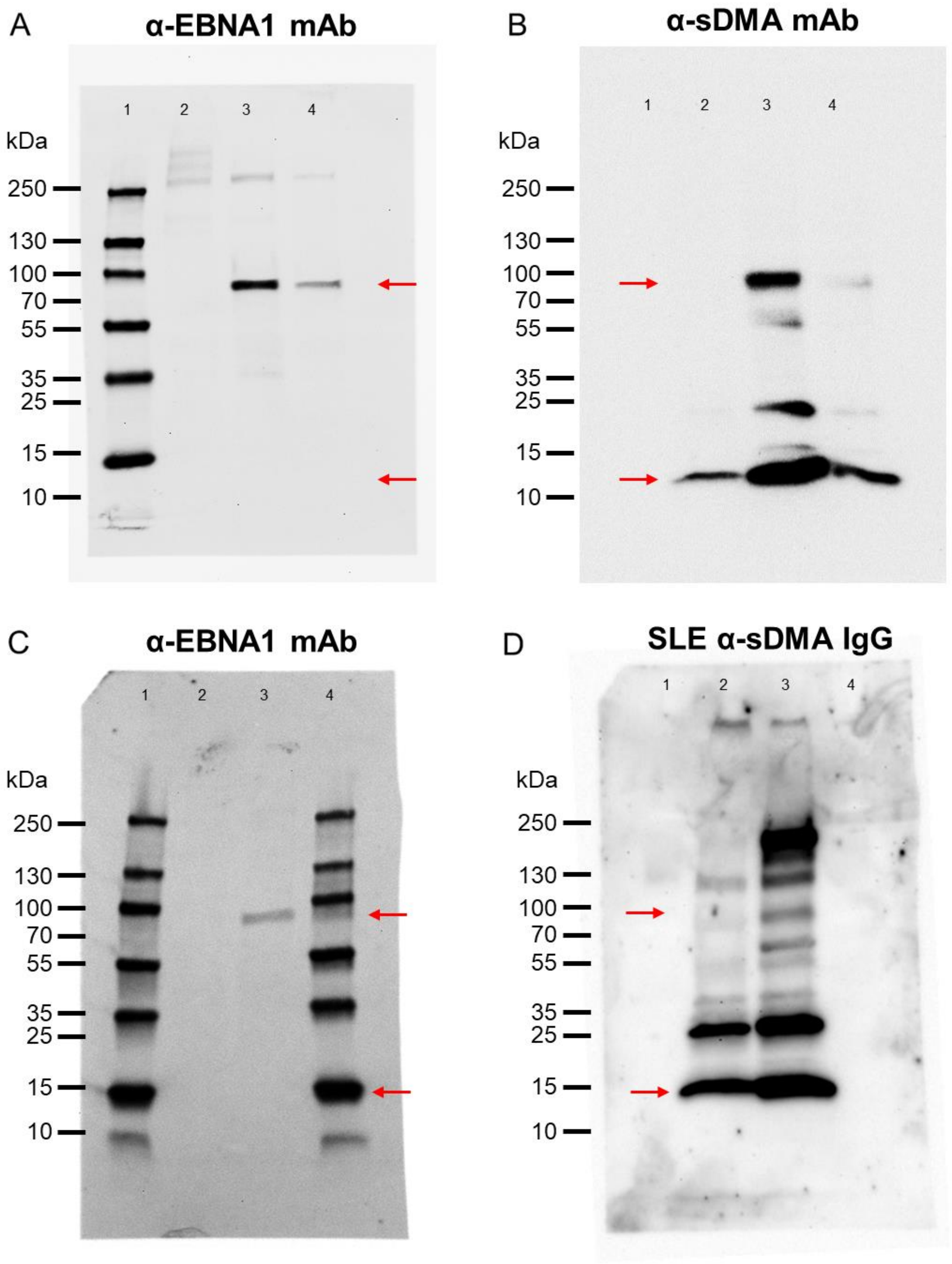


**Supplementary Figure 6: Uncropped and unedited blots for figure 3H/I.** Panels A and B correspond to Figure 3H; panels C and D correspond to Figure 3I. Arrows indicate the positions where the blots were cropped for the main figure. Lanes: 1, molecular weight ladder; 2, EBV- IMM B cell lysate; 3, EBV+ JY B cell lysate; 4, EBV+ DR7 B cell lysate. SLE, Systemic Lupus Erythematosus; sDMA, symmetrical dimethylarginine; mAb, monoclonal antibody; EBNA1, Epstein–Barr nuclear antigen 1; EBV, Epstein–Barr virus; IMM, BCL-6/Bcl-xL transduced immortalized; kDa, kilodalton.
